# Synergistic block of SARS-CoV-2 infection by combined drug inhibition of the host entry factors PIKfyve kinase and TMPRSS2 protease

**DOI:** 10.1101/2021.06.01.446623

**Authors:** Alex J.B. Kreutzberger, Anwesha Sanyal, Ravi Ojha, Jesse D. Pyle, Olli Vapalahti, Giuseppe Balistreri, Tom Kirchhausen

## Abstract

Repurposing FDA-approved inhibitors able to prevent infection by severe acute respiratory syndrome coronavirus 2 (SARS-CoV-2) could provide a rapid path to establish new therapeutic options to mitigate the effects of coronavirus disease 2019 (COVID-19). Proteolytic cleavages of the spike S protein of SARS-CoV-2, mediated by the host cell proteases cathepsin and TMPRSS2, alone or in combination, are key early activation steps required for efficient infection. The PIKfyve kinase inhibitor apilimod interferes with late endosomal viral traffic, and through an ill-defined mechanism prevents *in vitro* infection through late endosomes mediated by cathepsin. Similarly, inhibition of TMPRSS2 protease activity by camostat mesylate or nafamostat mesylate prevents infection mediated by the TMPRSS2-dependent and cathepsin-independent pathway. Here, we combined the use of apilimod with camostat mesylate or nafamostat mesylate and found an unexpected ~5-10-fold increase in their effectiveness to prevent SARS-CoV-2 infection in different cell types. Comparable synergism was observed using both, a chimeric vesicular stomatitis virus (VSV) containing S of SARS-CoV-2 (VSV-SARS-CoV-2) and SARS-CoV-2 virus. The substantial ~5-fold or more decrease of half maximal effective concentrations (EC_50_ values) suggests a plausible treatment strategy based on the combined use of these inhibitors.

**IMPORTANCE:** Infection by severe acute respiratory syndrome coronavirus 2 (SARS-CoV-2) is causing the coronavirus disease 2019 (COVID-2019) global pandemic. There are ongoing efforts to uncover effective antiviral agents that could mitigate the severity of the disease by controlling the ensuing viral replication. Promising candidates include small molecules that inhibit the enzymatic activities of host proteins, thus preventing SARS-CoV-2 entry and infection. They include Apilimod, an inhibitor of PIKfyve kinase and camostat mesylate and nafamostat mesylate, inhibitors of TMPRSS2 protease. Our research is significant for having uncovered an unexpected synergism in the effective inhibitory activity of apilimod used together with camostat mesylate or with nafamostat mesylate.

## INTRODUCTION

Severe acute respiratory syndrome coronavirus 2 (SARS-CoV-2) infection has caused the global pandemic known as coronavirus disease 2019 (COVID-2019). Currently there is no widespread use of an antiviral agent against the disease, but several candidates have been identified (1–6). Apilimod is currently in a clinical trial for the prevention of SARS-CoV-2 infections in the United States of America (ClinicalTrails.gov Identifier: NCT04446377). The TMPRSS2 protease inhibitor camostat mesylate has been tested with hospitalized COVID-19 patients in Denmark (7) and is in a clinical trial on adult COVID-19 patients in France (ClinicalTrails.gov Identifier: NCT04608266), while the higher affinity TMPRSS2 protease inhibitor, nafamostat mesylate, is being used in COVID-19 related trials in Russia (ClinicalTrials.gov Identifier: NCT04623021), South Korea (ClinicalTrails.gov Identifier: NCT04418128) and Japan (8).

Targeting the entry route of SARS-CoV-2 has been particularly challenging because there appear to be at least two different pathways for virus entry into cells (1). SARS-CoV-2 entry is mediated by the virus spike protein (9–11) and requires the receptor ACE2 in the host cell (1, 12, 13). Upon engagement of the ACE2 receptor, the spike protein catalyzes fusion of the viral membrane envelope with a host cell membrane to release the contents of the virus into the cytoplasm of host cells. For the spike protein to facilitate this reaction, it must first be cleaved by a host cell protease (1, 10). This can be accomplished by different host proteases including TMPRSS2 and TMPRSS4 (14), Factor Xa (15, 16), and by cathepsins during endocytosis (1). The protease inhibitors E-64 and camostat mesylate target cathepsin and TMPRSS2 and inhibit SARS-CoV-2 infection (1), but their low (micromolar) affinities have made them unlikely candidates for clinical use. Nafamostat mesylate inhibits TMPRSS2 with a high (nanomolar) affinity (2, 6). Apilimod has been shown to function in the endosomal pathway by inhibiting PIKfyve kinase causing a defect in viral trafficking prior to entry (3) but how this relates to the cathepsin or TMPRSS2 protease dependent pathways has not been investigated.

Using a chimeric vesicular stomatitis virus (VSV) in which the attachment and fusion glycoprotein G is replaced by the spike S protein of SARS-CoV-2 (VSV-SARS-CoV-2) (17) we investigate inhibition of infection in different cell types known to contain different levels of cathepsin and TMPRSS2 proteases. We tested combinations of the protease inhibitors E-64 (cathepsin), camostat mesylate and nafamostat mesylate (TMPRSS2) and the lipid kinase inhibitor apilimod (PIKfyve kinase) on VSV-SARS-CoV-2 infection of multiple cell types. We observed a 5-fold synergistic effect in the infection of VSV-SARS-CoV-2 by simultaneous inhibition of TMPRSS2 and PIKfyve kinase. Furthermore, the synergistic inhibitory effects on infection by VSV-SARS-CoV-2 or a clinical isolate of SARS-CoV-2 were observed with nanomolar concentrations of camostat mesylate and apilimod. This finding suggests that a combination of inhibitors that target two different host factors in the entry pathway of SARS-CoV-2 will likely be more effective than targeting either alone.

## RESULTS AND DISCUSSION

### VSV-SARS2-CoV-2 infection is partially prevented by PIKfyve kinase or TMPRSS2 protease inhibitors

It has been proposed that two distinct host proteases, TMPRSS2 and cathepsin facilitate different SARS2-CoV-2 viral entry routes - cell surface or endosomal - and their abundance in different cell types may influence the entry pathway (1). To test this, we employed a panel of inhibitors on SARS-CoV-2 infection in African green monkey kidney epithelial derived VeroE6 cells poorly expressing TMPRSS2, VeroE6 cells stably expressing ectopic TMPRSS2, as well as human colon carcinoma derived Caco-2 cells and human lung derived Calu-3 cells naturally expressing TMPRSS2.

We first established the relative importance of the cathepsin-dependent route for infection of VSV-SARS-CoV-2 by determining the effect of the cathepsin inhibitor E-64 on expression of eGFP mediated by VSV-SARS-CoV-2. As summarized in the plots in Fig. 1A, we found a cell type dependence in the extent of infection block with a half maximal effective concentration (EC_50_) in the ~5-10 μM range. Notably, only Vero E6 cells treated with E-64 displayed a full infection block, the result expected for cells expressing cathepsin but no TMPRSS2; in contrast, cells expressing cathepsin and TMPRSS2 (VeroE6 + TMPRRS2, Caco-2) displayed a partial block, while Calu-3 cells with minimal expression of cathepsin L (18) did not respond, as expected, to E-64 treatment (1).

**Fig. 1.**
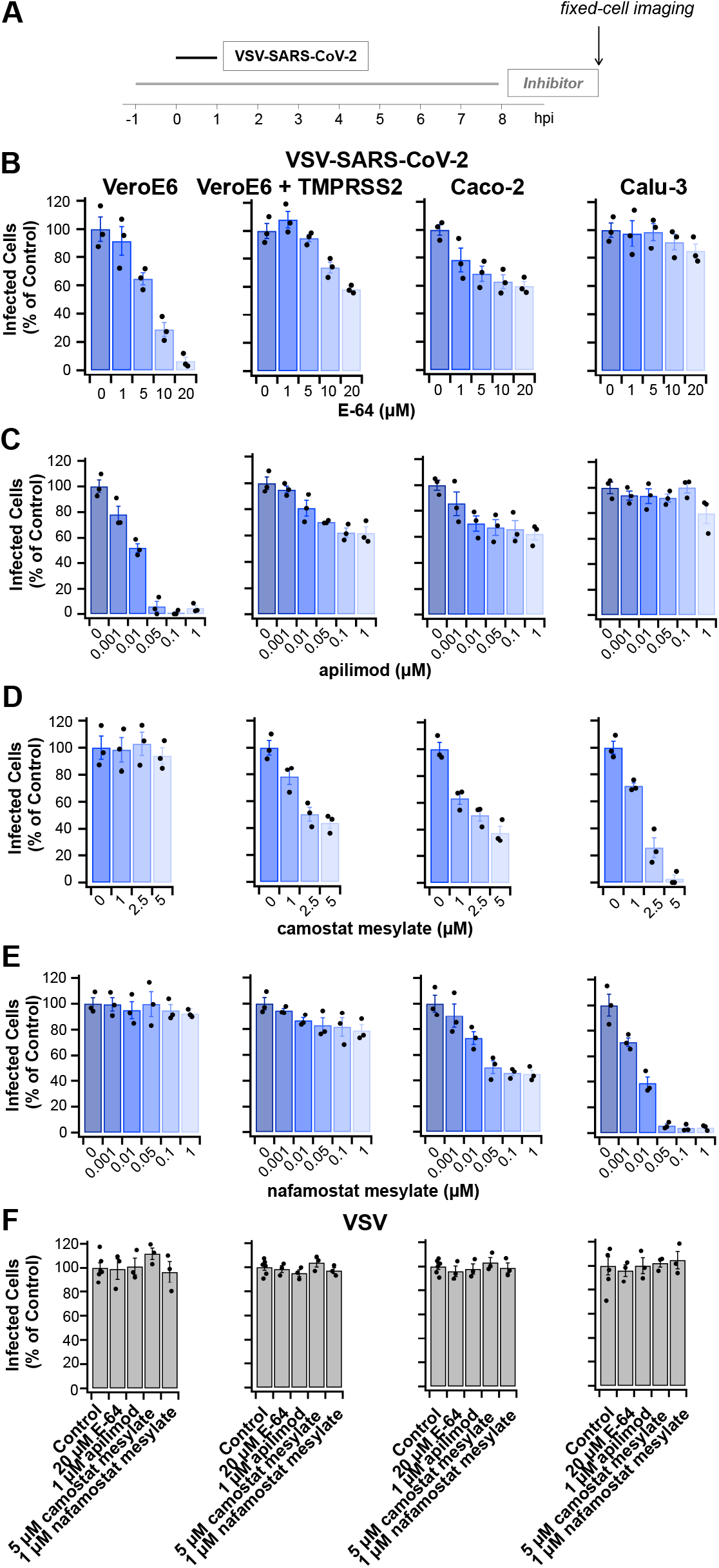
Protease inhibitors E-64, apilimod, camostat mesylate or nafamostat mesylate prevent infection by VSV-SARS-CoV-2 but not by VSV. **(A)** Schematic of infectivity assay for cells pretreated for 1 h or not with the inhibitors, subsequently infected with VSV-SARS-CoV-2 for 1 h in the presence or absence of inhibitors. The cells were incubated for another 7 h in the presence or absence of inhibitors and then fixed; the percentage of cells expressing eGFP measured by spinning disc confocal microscopy. **(B-F)** Quantification of the number of infected cells from three independent experiments, each determined from 5 fields of view containing 80-200 cells per experiment (error bars show SEM) for the indicated cell types. Infected Vero **(B)** or Vero + TMPRSS2 cells **C**) were analyzed 8 hpi using 0.5 μg/mL VSV-SARS-CoV-2 RNA. Infected Caco-2 **(D)** or Calu-3 cells **(E**) were analyzed 8 hpi using 5 μg/mL VSV-SARS-CoV-2 RNA. Cells infected with 0.075 μg/mL VSV RNA **(F)** were analyzed 6 hpi. In each case, these virus concentrations and conditions of infection corresponded to an MOI of ~ 0.5.

We extended the analysis and studied the inhibitory effect by the PIKfyve kinase inhibitor apilimod on VSV-SARS-CoV-2 infection (Fig. 1 B). We found that all cell types shown to be sensitive to E-64 also responded to treatment with apilimod with EC_50_ ~ 10 nM whereas Calu-3, which is insensitive to E-64 (19), did not respond. Although apilimod doesn’t inhibit cathepsin B or L (20), these observations are consistent with potential modulation of the endosomal availability of cathepsin by the activity of PIKfyve kinase (21, 22).

A similar analysis, to document the reduction of viral infection using camostat mesylate to block the protease activity of TPMRSS2 (Fig. 1C) or nafamostat mesylate (Fig. 1D) showed, as anticipated, no response in Vero E6 cells lacking TMPRSS2, partial inhibition in Vero E6 or Caco-2 cells expressing TMPRSS2 or full inhibition in Calu-3 cells naturally expressing TMPRSS2 but insensitive to cathepsin inhibition.

These observations are consistent with previous results (1, 2, 6, 19) that infection by VSV-SARS-CoV-2 occurred through two complementary entry pathways with different importance depending on the cell type, one depending on the proteolytic activity of cathepsin and the second relying of the proteolytic activity of TMPRSS2. As a negative control for these experiments, we included infection by the parental VSV (Fig. 1E), whose ability to infect host cells is known to be independent of the enzymatic activities of cathepsin, TMPRSS2, or PIKfyve kinase (1–4, 6). As expected, none of the inhibitors for these enzymes influenced the extent of VSV infectivity in any of the cells used. From these results we could also exclude potential cytotoxic effects by the compounds in the concentration range used.

### Synergistic prevention of VSV-SARS2-CoV-2 infection by combined use of PIKfyve kinase and TMPRSS2 protease inhibitors

Since the cathepsin and TMPRSS2-dependent activation of SARS-CoV-2 correspond to complementary entry pathways thought to act independent of each other, we expected an additive inhibitory effect upon their simultaneous inhibition in cells that express both proteases. Indeed, VSV-CoV-2 infection of Vero E6 cells expressing cathepsin and TMPRSS2 (VeroE6 +TMPRSS2) were equally inhibited (EC_50_ ~ 2 μM) by camostat mesylate in the absence or increasing concentrations of E-64 up to 20 μM, the concentration at which E-64 maximally blocked VSV-CoV-2 infection (Fig. 2A). We used SynergyFinder 2.0 (23) to compare the combined response obtained experimentally with the expected outcome calculated by the Bliss synergy-scoring model. Using this reference model, that considers multiplicative effect of single drugs as if they acted independently (23, 24), we obtained an overall δ-score of 6 indicative of an additive interaction between camostat and E-64. At combined high levels of camostat and E64, these protease inhibitors also showed weak concentration dependence, consistent with the prediction of a published model (25). In contrast, combined use of camostat mesylate and apilimod led to enhanced inhibition of VSV-CoV-2 infection with a ~ 2-fold decrease in the EC_50_ of camostat mesylate (from 1 to 0.4 μM) (Fig. 2B), suggestive of a synergy effect.

**Fig. 2.**
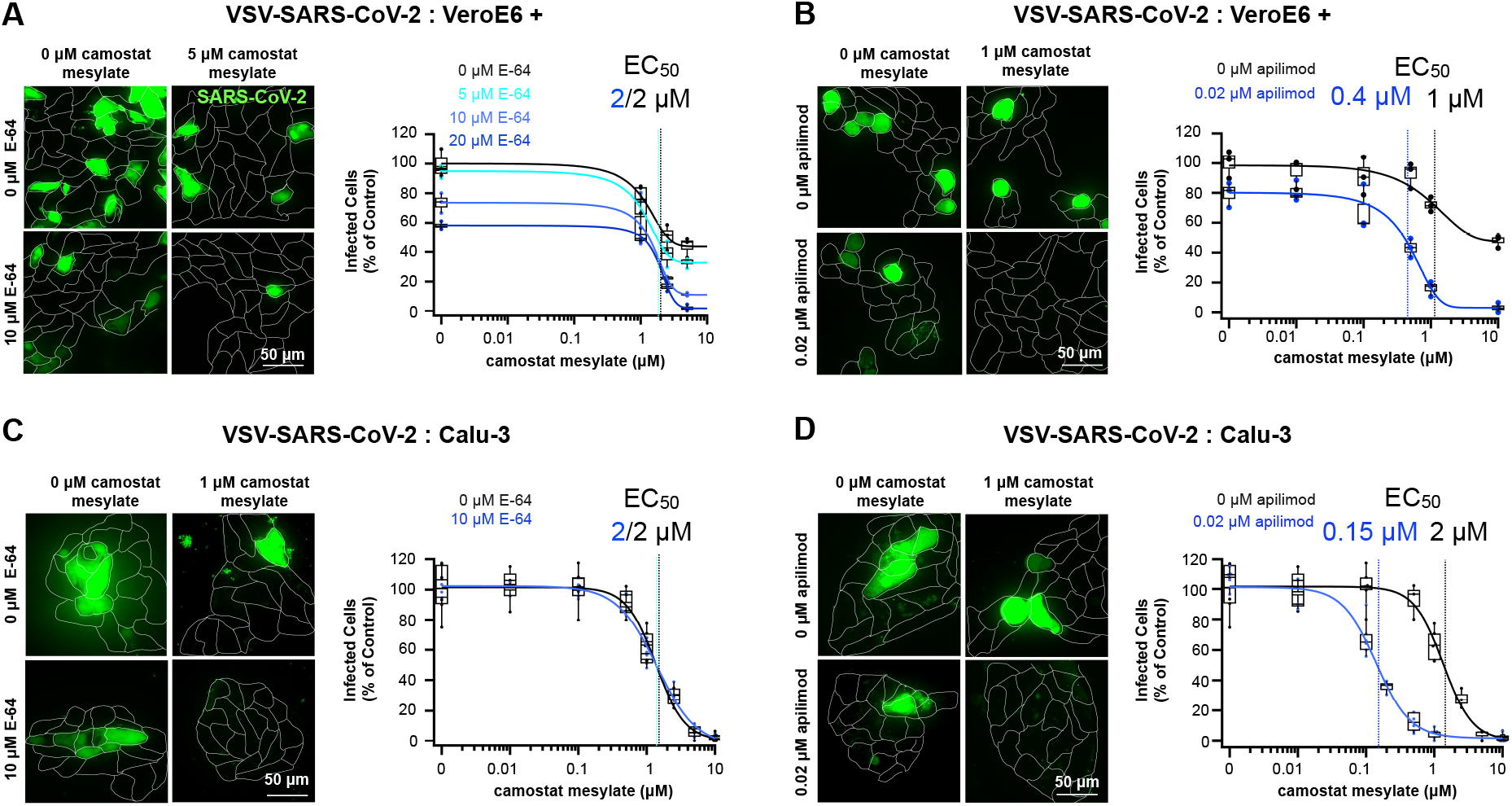
Synergistic inhibition of VSV-SARS2-CoV-2 infection by combined use of apilimod and camostat mesylate. Data are from infection results using VSV-SARS2-CoV-2 obtained with different cell types in the absence or combined presence at increasing concentrations of E-64 and camostat mesylate **(A, C)** or apilimod and camostat mesylate **(B, D)**. Representative maximum-Z projections views (left panels) are from whole-cell volume images obtained with optical sections separated by 0.5 μm using spinning disk confocal microscopy; cells were infected with 0.5 μg/mL viral RNA VSV-SARS-CoV-2 and imaged 8 hpi. Scale bar: 50 μm. Corresponding quantifications of infection (right panels) are shown in the plots. Each point corresponds to one independent experiment; the data represent results from 5 fields of view containing 80-200 cells per experiment. Estimated EC_50_’s are indicated. Significance in the difference of EC_50_ was determined from fitting replicated experiments and using an unpaired t-test; P-values were 0.02 for **(B)** and 0.002 for **(D);** there was no statistically significant difference in the EC_50_ values for the experiments reported in **(A)** or **(C)**.

We carried a similar set of experiments using Calu-3 cells known to be deficient in Cathepsin-L (18, 26) and which are poorly susceptible to infection of filoviruses mediated by the cathepsin-dependent infection route (18). These cells are insensitive to inhibition by of VSV-SARS-CoV-2 and SARS-CoV-2 infection by E-64 (1, 19, 27, 28) (Fig.1B) and apilimod ((19, 29), Fig. 1C). While presence of E-64 did not affect the inhibition profile of camostat mesylate (Fig. 2C), we detected a ~ 5-fold decrease from 1 μM to 0.2 μM in the EC_50_ of apilimod in cells simultaneously treated with 10 μM E-64 (Fig. 2D). This result was unexpected, given that infection of the Calu-3 cells by SARS-CoV-2 didn’t appear to be affected by apilimod ((29) and Fig. 1C).

We extended the synergy analysis to verify the effects upon simultaneous use of variable amounts of apilimod and nafamostat mesylate, another inhibitor of TMPRSS2. Our data also indicates enhanced infection inhibition for nafamostat mesylate with a ~ 20-fold decrease of EC_50_ from 0.02 to 0.001 μM (Fig. 3A, central panel). Simultaneous treatment with apilimod and increasing amounts of nafamostat mesylate led to an analogous enhanced potency of apilimod inhibition with a ~10-fold decrease of EC_50_ from 0.01 to 0.001 μM (Fig. 3A, right panel). Formal evaluation for synergism using the Bliss reference model was consistent with a synergy δ-score of 41 (a score greater than 10 indicates synergy (23)).

**Fig. 3.**
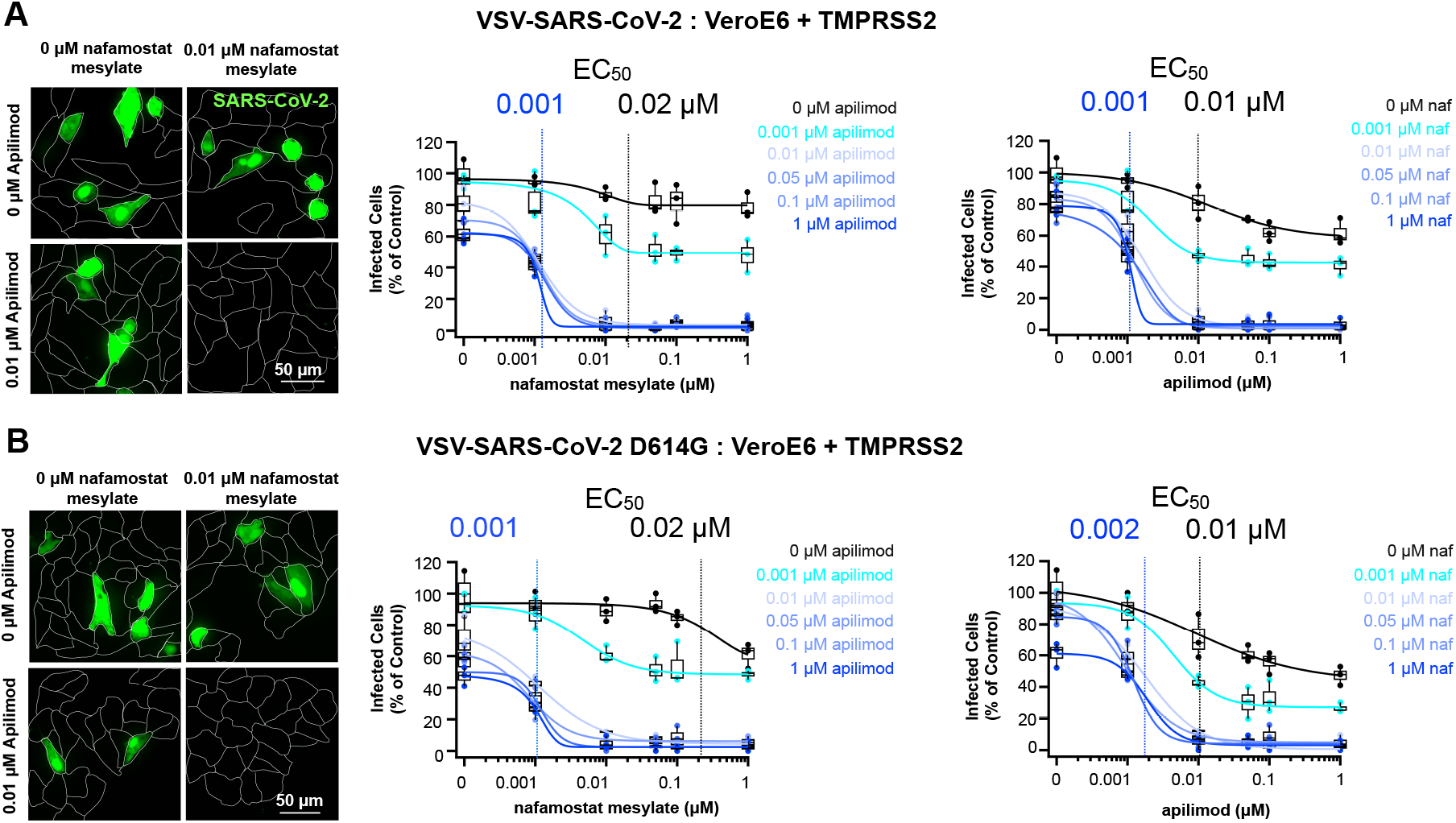
Synergistic inhibition of VSV-SARS2-CoV-2 infection by combined use of apilimod and nafamostat mesylate. Data are from infection results obtained with VeroE6 + TPRSS2 cells in the absence or combined presence at increasing concentrations of apilimod and nafamostat mesylate. Representative maximum-Z projections views (left panels) are from whole-cell volume images obtained with optical sections separated by 0.5 μm using spinning disk confocal microscopy; cells were infected with 0.5 μg/mL viral RNA VSV-SARS-CoV-2 **(A)** or 0.2 μg/mL VSV-SARS-CoV-2 D614G viral RNA **(B)** and imaged 8 hpi. Scale bar: 50 μm. Corresponding quantifications of infection (right panels) are shown in the plots. Each point corresponds to one independent experiment; the data represent results from 5 fields of view containing 80-200 cells per experiment. Estimated EC_50_’s are indicated. Both plots in the central and right panels used the same data.

Comparable strong synergistic effects (δ-score of 34) by combined use of apilimod and nafamostat mesylate were observed in Vero E6 cells expressing TMPRSS2 (Fig. 3B, central and right panels) infected with VSV-SARS-CoV-2 D614G whose spike S protein includes the point mutation known to increase infectivity of the native SARS-CoV-2 (30). While this mutation has no effect on the virus entry mechanism or sensitivity to proteases, it stabilizes the stalk region of the spike, which otherwise tend to fall apart after furin cleavage between S1 and S2 (30–33).

Taken together, these synergy results highlight the unexpected non-additive involvement of PIKfyve kinase activity for the functional effectiveness of TMPRSS2 to mediate viral entry along the TMPRSS2 route.

### Synergistic prevention of SARS2-CoV-2 infection by combined use of PIKfyve kinase and TMPRSS2 protease inhibitors

To determine whether SARS-CoV-2 virus also display an enhanced block of infection upon combined use of apilimod and camostat mesylate we infected Caco-2 cells ectopically expressing hACE2, the main receptor for SARS-CoV-2. This cell line was created as a way to enhance the susceptibility of Caco-2 to infection by SARS-CoV-2 since the parental cells express low endogenous levels of ACE2 (34); successful infection was scored 18 hrs post infection by the appearance of viral N protein using immunofluorescence microscopy (Fig. 4, left panel). The inhibitory EC_50_ of camostat mesylate decreased ~ 6-fold from 0.6 to 0.1 μM when used together with apilimod, (Fig. 4, right panel). This observation extends our results from VSV-SARS-CoV-2 chimeras to SARS-CoV-2 virus, and illustrates that combined chemical inhibition of PIKfyve kinase and TMPRSS2 protease activities is likely to also prevent SARS-CoV-2 infection with strong synergy.

**Fig. 4.**
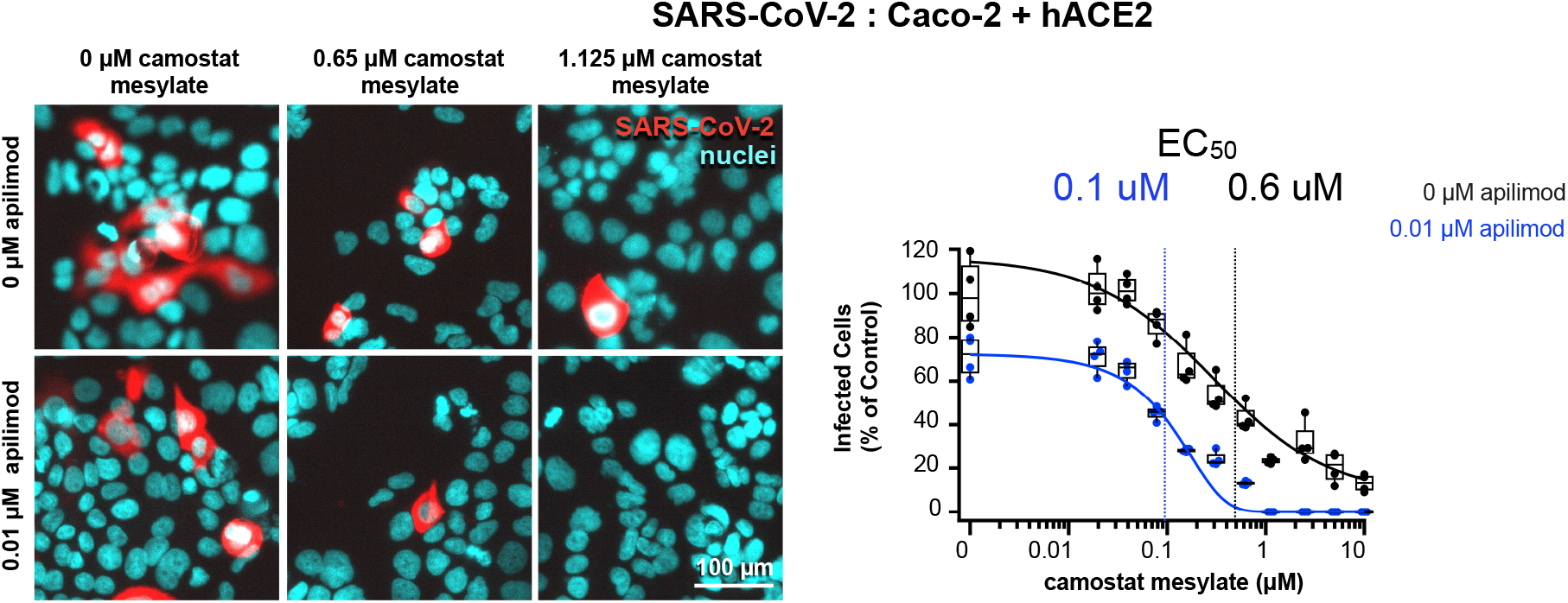
Synergistic inhibition of SARS2-CoV-2 infection by combined use of apilimod and camostat mesylate. Natural SARS-CoV-2 infection inhibition in Caco-2 + hACE2 cells (MOI = 0.5) by camostat mesylate in the absence or combined presence of apilimod. Representative examples of images of fixed samples stained with an antibody specific for N (red) and with Hoechst DNA stain (cyan) to identify the nuclei obtained using wide field epifluorescence microscopy (left panels) and corresponding quantifications of infection (right panels) are shown. Each data points were from 4 independent experiments, representing results from 6,000-10,000 cells per experiment. Estimated EC_50_’s are shown. Significance in the difference (P-value of 0.01) of EC_50_ was determined using an unpaired t-test after a least square non-linear fitting of the data curves from replicated experiments.

### Final remarks

According to our present understanding of early steps necessary for successful infection, SARS-CoV-2 would enter cells using two main redundant routes, each requiring cleavage of the viral S spike protein, one dependent on the proteolytic activities of members of the cathepsin family, the other requiring proteolysis by TMPRSS2 or similar transmembrane-serine proteases. Expression level of these proteases depends on cell type and hence their relative importance to support successful infection would hinge in part on their relative expression. Cathepsin is primarily found in late endosomes or lysosomes and requires low pH for optimal enzymatic activity; for these reasons it has been proposed that SARS-CoV-2 entry occurs from the endolysosomal compartment (35, 36). In contrast, TMPRSS2 is thought to be at the cell surface (37) and its optimal proteolytic activity is pH independent (38); hence it has been inferred that TMPRSS2 cleavage of spike S protein occurs on virions at the plasma membrane from which viral entry is then assumed to occur (1).

Not surprising for protease inhibitors with different targets, combined inhibition of the enzymatic activities of cathepsin and TMPRSS2 by the respective protease inhibitors E-64 and camostat mesylate led to additive prevention of infection by SARS-CoV-2 (1) and by the VSV-SARS-CoV-2 chimera (this study).

Apilimod specifically inhibits PIKfyve kinase thereby blocking accumulation of PI(3,5)P2, a key phosphoinositide required to modulate the function of a number of proteins involved in late endosomal traffic (3, 21, 39, 40). While it is now well established that interference with PIKfyve kinase prevents infection of a selected group of viruses including Ebola (20), VSV-Ebola (3), Marburg (20), VSV-SARS-CoV-2 (3) and native SARS-CoV-2 (3–5), the molecular mechanism events responsible for the interference remain to be determined. It has been shown that Apilimod has anti-Ebola infection synergy with clomiphene cytrate (δ-score of 23.8; re-calculated from data in Fig. 1B of (41) using the Bliss reference model (23)) and also with other drugs that interfere, by unknown mechanisms, with Ebola entry in late endosomes (41). We have now shown synergistic inhibition of SARS-CoV-2 infection by combined use of apilimod and camostat or apilimod and nafamostat mesylate, even in cells as diverse as Vero (which have an active cathepsin entry pathway) and Calu-3 (which lack it).

Prevailing models for the cellular location of TMPRSS2-activated SARS-CoV-2 entry favor fusion at the cell surface rather than from endosomal compartments, because TMPRSS2 activity is pH independent. Ebola or SARS-CoV-2 viruses accumulate in EEA1 containing early endosomes (3, 20), a behavior consistent with the disruption of endolysosomal traffic that occurs when PIKfyve kinase activity is inhibited by genetic or pharmacological means (21, 40, 42). It is therefore possible that changes in the endosomal milieu, combined with redirection of SARS-CoV-2 and/or TMPRSS2 traffic in the presence of apilimod, might make infection more dependent on TMPRSS2 pharmacological inactivation than in its absence. Combined presence of apilimod and camostat or nafamostat mesylate would manifest as synergy of these compounds to prevent infection. These observations would also be consistent with favored entry of TMPRSS2-activated SARS-CoV-2 from internal membrane compartments rather than from the cell surface.

The strong synergy we described in the laboratory setting for single cell types upon combined use of the PIKfyve kinase and TMPRSS2 inhibitors highlights the potential clinical advantage of using a similar strategy to ameliorate the viral load and potentially reduce the risk to COVID-19 in patients infected with SARS-CoV-2 based on simultaneous inhibition of these enzymes. Variants of apilimod and camostat, rendered more soluble by mesylate modification, are currently in clinical trials as orally administered inhibitors dispensed alone for various indications, including COVID-19 (7, 43, 44). More soluble nafamostat mesylate has been administered intravenously to treat COVID-19 patients (8). We do not yet know how efficiently these inhibitors reach nasopharyngeal and lung tissues; nevertheless, we surmise that combined use of these PIKfyve and TMPRSS2 inhibitors might also recapitulate the ~5-10-fold increase in inhibitor efficiency that we observe in the laboratory setting, offering considerable advantage for clinical application.

## MATERIAL AND METHODS

### Materials

These reagents were purchased as indicated: Dulbecco modified Eagles Medium (DMEM) supplemented with 4.5 g/L glucose, L-glutamine, and sodium pyruvate (Corning, Inc., cat. 10-013-CV), fetal bovine serum (FBS, Atlanta Biologicals, cat. S11150H), penicillin-streptomycin 100x solution (VWR, cat. 45000-652), camostat mesylate (Sigma-Aldrich, cat. SML0057), E-64 (Santa Cruz Biotechnology, cat. sc-201276), nafamostat mesylate (Cayman Chemical Company, cat. #14837), apilimod (MedChem Express, cat. HY-14644), dimethyl sulfoxide (DMSO, Sigma-Aldrich, cat. 26855), Round cover glass #1.5, 25 mm (Cellvis cat. C8-1.5H-N), Polydimethylsiloxane (PDMS, SYLGARD from Krayden, cat. DC4019862), isopropyl alcohol (VWR, cat. 9080-03/MK303108), potassium hydroxide (Sigma-Aldrich, cat. 484016), wheat germ agglutinin (WGA)-Alexa647 (Invitrogen, cat. W32466), paraformaldehyde (Sigma-Aldrich, cat. P6148), sodium phosphate dibasic (Thermo Fisher Scientific, cat. BP329-1), potassium phosphate monobasic (Sigma-Aldrich, cat. P5379), sodium chloride (EMD Millipore, cat. SX0420-5), and potassium chloride (Thermo Fisher Scientific, cat. P217-500), TRIS (Goldbio, cat. T-400-500), EDTA (Sigma-Aldrich, cat. E5134), sucrose (Sigma-Aldrich, cat. S0389), fetal calf serum (FCS, Hyclone, cat. SH30073.03), bovine serum albumin (GE Healthcare), Triton-X (Thermo Fischer, cat. 28314), Hoechst DNA stain (Thermo Fischer, cat. 62249), L-glutamine (Sigma, cat. G7513), Hoechst DNA dye (cat. H6021, Sigma-Aldrich), and Alexa 647 fluorescently labelled goat anti-rabbit antibody (Thermo Fisher, cat. A32733).

### Purification of VSV-SARS-CoV-2 chimeras

Generation of recombinant VSV (Indiana serotype) expressing eGFP (VSV-eGFP) and VSV-eGFP chimeras whose glycoprotein G was replaced with either the wild type spike S protein of SARS-CoV-2 Wuhan-Hu-1 strain (VSV-SARS-CoV-2) or with the point mutant D614G (VSV-SARS-CoV-2 D614G) was done as described (17). Briefly, VSV was grown by infection of BSRT7/5 and VSV-SARS-CoV-2 chimeras were grown by infection of MA104 cells. Cells were grown in 12-18 150 mm dishes and infected at an MOI of 0.01. Virus-containing supernatant was collected at 48 hours post infection. The supernatant was clarified by low-speed centrifugation at 1,000 × g for 10 min at room temp. An initial pellet of virus and extracellular particles was generated by centrifugation in a Ti45 fixed-angle rotor at 30,000 × g (25,000 rpm) for 2 hours at 4°C. The pellet was resuspended overnight in 1X NTE (100 mM NaCl, 10 mM Tris-HCl pH 7.4, 1 mM EDTA) at 4°C. The resuspended pellet was layered on top of a 15% sucrose-NTE cushion and subjected to ultracentrifugation in a SW55 swinging bucket rotor at 110,000 × g (35,000 rpm) for 2 hours at 4°C. The supernatant was aspirated and virus pellet resuspended in 1X NTE overnight at 4°C. The resuspended virus pellet was separated on a 15-45% sucrose-NTE gradient by ultracentrifugation in a SW55 swinging bucket rotor at 150,000 × g (40,000 rpm) for 1.5 hours at 4°C. The predominant light-scattering virus band was observed in the lower third of the gradient and was extracted by side puncture of the gradient tube. Extracted virus was diluted in 1X NTE and collected by ultracentrifugation in a Ti60 fixed-angle rotor at 115,000 × g (40,000 rpm) for 2 hours at 4°C. The final pellet was re-suspended overnight in 1X NTE in a volume of 0.2 to 0.5 mL depending on the size of the pellet and stored at 4°C for use in subsequent imaging experiments.

### Isolation and propagation of SARS-CoV-2

Human samples were obtained under the Helsinki University Hospital laboratory research permit 30 HUS/32/2018 § 16. Isolation of SARS-CoV-2 from a COVID-19 Briefly, a nasopharyngeal swab in 500 μl of Copan UTM® Universal Transport Medium was inoculated on Calu-3 cells (P1) and incubated for 1 h at 37°C, after which the inoculum was removed and replaced with Minimum Essential Medium supplemented with 2% FBS, L-glutamine, penicillin and streptomycin. Virus replication was determined by RT-PCR for SARS-CoV-2 RdRP (45), and the infectious virus collected 48h after inoculation. The P1 stock was propagated once (P2) in VeroE6+TMPRSS2, sequenced and stored at −80 °C. Virus stocks were stored in DMEM, 2% FCS, 2 mM L-glutamine, 1% penicillin-streptomycin.

### Cell Culture

VeroE6 (ATCC CRL-1586), Caco-2 (ATCC HTB-37), and Calu-3 cells (ATCC HTB-55) were purchased from ATTC. VeroE6+TMPRSS2 cells were a gift from Siyan Ding (14). VeroE6, VeroE6+TMPRSS2, Caco-2, and Calu-3 cells were maintained in DMEM supplemented with 25 mM HEPES, pH 7.4, 10% fetal bovine serum and 1% penicillin-streptomycin. VeroE6 and VeroE6+TMPRSS2 cells were grown at 37°C and 5% CO2 and split at a ratio of 1:10 every 3-4 days when cells were at ~90% confluency. Caco-2 cells were grown at 37°C and 5% CO2 and split at a ratio of 1:5 every 3-4 days when cells were at ~95% confluency. Calu-3 cells were grown at 37°C and 7% CO2 and split at a ratio of 1:3 every 5-6 days when cells were at ~95% confluency. BSR-T7 cells were derived from BHC cells (46) and grown in DMEM supplemented with 10% FBS and 1% penicillin-streptomycin. BSR-T7 were grown at 37°C and 5% CO2 and split at a ratio of 1:20 every 2-3 days when cells were at ~90% confluency. MA104 cells (ATCC, CRL-2378.1) were grown in Media 199 supplemented with 10% FBS and 1% penicillin-streptomycin. MA104 cells were grown at 37°C and 5% CO2 and split at a ratio of 1:3 every 2 days when cells were at ~90% confluency. The media was changed in all cell types every 2 days and regularly tested for presence of mycoplasma.

Caco-2 cells stably expressing human ACE2 were generated by transduction with third generation lentivirus pLenti7.3/V5 DEST ACE2-EmGFP (prepared by the cloning facility Dream-Lab, Institute of Biotechnology, University of Helsinki, Helsinki, Finland; the expression of Emerald GFP is driven by the SV40 promoter. Cells expressing eGFP were isolated by FACS. These cells were grown in DMEM media supplemented with 10% fetal calf serum, 1% penicillin-streptomycin, and 2 mM L-glutamine. Cells were kept at 37°C, 5% CO2, and split every 2-4 days when ~90% confluent.

### Infection protocol for VSV and VSV-SARS-CoV-2

Polydimethylsiloxane was cured by vigorously mixing with curing reagent at a ratio of 1 to 10. PDMS was poured into 10 cm petri dish (5 g of PDMS per plate) and incubated at 90 °C for ~4 hours. Plates were removed and stored at room temperature until use. PDMS was removed from petri dish using a razor blade. 3 mm holes were punched into PDMS which was cut with a blade to be ~6-10 mm. Glass slides were cleaned by sonication first in isopropanol for 20 min then in 0.5M potassium hydroxide for 20 min followed by extensive washing in Milli-q water. Glass was dried in an oven 60 °C for 30 minutes then bonded to the PDMS by exposing PDMS and glass to air plasma at 750 mTorr, 30 W for 2 minutes using a PDC-001 plasma cleaner (Harrick Plasma) then firmly pressing glass and PDMS together. This was followed by placing bonded glass and PDMS in an oven at 90 °C for 20 minutes. The glass cover slips mounted with a PDMS well were then placed in 70% ethanol for 10 minutes to sterilize prior to use for cell culture.

The day prior to the experiment cells were plated in PDMS wells on the glass slide stored in a 6 well plate at a density to achieve ~70% confluence the day of the experiment. On the day of the experiment cells were incubated with the desired inhibitor concentration for 1 hour. Media was then removed and virus that had been diluted into media containing the indicated inhibitor concentration was added to the well in a volume of 10 μL. Media was left in the 6 well plate outside of the PDMS well at a level less than the height of the PDMS well to maintain humidity and prevent evaporation. After the virus was incubated with the cells for 1 hour the cells were washed with media containing the indicated inhibitor and then the well was filled with fresh media. In all experiment’s cells were kept at 37°C and 5% (for Vero, Vero+TMRPSS2, and Caco-2 cells) or 7% CO2 (for Calu-3 cells) and media was pre warmed to 37°C. At 6 hours after initiating the infection with VSV or 8 hours after initiating the infection with VSV-SARS-CoV-2 media was removed and the cells were stained by adding 5μg/mL WGA-Alexa647 in PBS (137 mM NaCl, 2.7 mM KCl, 8 mM Na_2_HPO_4_, 2 mM KH_2_PO_4_, pH 7.4) for 30 seconds at room temperature. Cells were then washed with sterile PBS 2 times and then fixed with 4% paraformaldehyde in PBS. Infected cells were imaged using a spinning disk confocal microscope with a 40X oil objective with a pixel size of 0.33 μm where 20 optical planes were taken at 0.5 μm apart for every field of view (47). Cells were considered infected when they displayed a cytosolic eGFP fluorescence signal with a relative intensity of 1.4 times that of the background of uninfected cells. Example images are max intensity projections of the cell volume where the outline (white) line was obtained by tracing the WGA-Alexa647 signal outlining the cell.

### Infection protocol for SARS-CoV-2

All experiments with SARS-CoV-2 were performed in BSL3 facilities at the University of Helsinki with appropriate institutional permits. Virus samples were obtained under the Helsinki University Hospital laboratory research permit 30 HUS/32/2018 § 16. Virus titers were determined by plaque assay in VeroE6+TMPRSS2 cells. Cells in DMEM, supplemented with 10% FBS, 2 mM L-glutamine, 1% penicillin-streptomycin, 20 mM HEPES (pH 7.2 were seeded 48 h before treatment at a density of 15,000 cells per well in 96-well imaging plates (PerkinElmer cat. 6005182). Inhibitors, or DMSO control, were either added 60 min prior to infection, or added 90 min post infection at a multiplicity of infection (MOI) 0.5 plaque forming units per cell. Infections were carried for 20 h at 37 °C and 5% CO2. Cells were then fixed with 4% paraformaldehyde in PBS for 30 min at room temperature before being processed for immuno-detection of viral N protein, automated fluorescence imaging and image analysis. Briefly, viral NP was detected with an in house developed rabbit polyclonal antibody (34) counterstained with Alexa-647-conjugated goat anti rabbit secondary antibody, and nuclear staining done using Hoechst DNA dye. Automated fluorescence imaging was done using an epifluorescence high-content Molecular Device Image-Xpress Nano microscope equipped with a 20x objective and a 4.7Mpixel CMOS camera (pixel size 0.332 μm). Image analysis was performed with the software CellProfiler-3 (www.cellprofiler.org). Automated detection of nuclei was obtained using the Otsu algorithm inbuilt in the software. To automatically identify infected cells, an area surrounding each nucleus (5 pixels expansion of the nuclear area) was used to estimate the fluorescence intensity of the viral NP immuno labeled protein, using an intensity threshold such that less than 0.01% of positive cells were detected in non-infected wells.

### Statistical analysis

The significance of response (synergy δ-score) upon combined use of two drugs (Fig. 3) was calculated using the Bliss reference model for combination of two drugs incorporated in the stand-alone web-application SynergyFind v2 (23). This model assumes a stochastic process in which the effects of two drugs act independently. δ-score values between −10 and 10 suggest an additive interaction; δ-scores larger than 10 suggest the interaction is synergistic.

The EC5_50_ values in Figures 2 B, C, and Figure 4 were obtained from replicate determinations calculated with least-square nonlinear curve fitting using Igor Pro (WaveMetrics). An unpaired T-test was then used to determine the statistical significance in the difference in the values of EC_50_ of camostat mesylate as a function of apilimod.

## AUTHOR CONTRIBUTIONS

We thank Sean Whelan for comments and suggestions. Alex J.B. Kreutzberger carried out all the experiments in Figs. 1-3. Ravi Ojha and Giuseppe Balistreri carried out the experiments in Fig. 4. Anwesha Sanyal maintained the cells lines and assisted with infectivity assays. Ravi Ojha generated Caco2-ACE2+EmGFP cells and propagated SARS-CoV-2 virus under the supervision of Olli Vapalahti. Jesse Pyle generated the stocks of VSV-SARS-CoV-2 chimeras. We thank Tegy John Vadakkan for maintaining the spinning disc confocal microscope. This research was supported by a NIH Maximizing Investigators’ Research Award (MIRA) GM130386, by research grants from the Danish Technical University and SANA and unrestricted funds from IONIS to T.K; Harvard Virology Program, NIH training Grant T32 AI07245 postdoctoral fellowship to A.J.B.K.; by an Academy of Finland research grant 336490 by the Jane and Aatos Erkko Foundation, by the EU Horizon 2020 program VEO 874735, and by Helsinki University Hospital Funds TYH2018322 to O.V; by a research grant from the Academy of Finland grant 318434 and private funds supporting COVID-19 research to G.B.; by the University of Helsinki graduate program in Microbiology and Biotechnology to R.O. Tom Kirchhausen and Alex J.B. Kreutzberger were responsible for the overall design of the study; Tom Kirchhausen and Alex Kreutzberger drafted the manuscript; all authors commented on the manuscript.

